# Maternal cradling bias drives early handedness in infant monkeys: A longitudinal study of grasping lateralization in baboons

**DOI:** 10.1101/2021.03.09.434410

**Authors:** Grégoire Boulinguez-Ambroise, Emmanuelle Pouydebat, Éloïse Disarbois, Adrien Meguerditchian

**Affiliations:** Laboratoire de Psychologie Cognitive UMR7290, CNRS, Aix-Marseille Univ, Institut Language, Communication and the Brain, France; Station de Primatologie UAR846, CNRS, Rousset-sur-Arc, France; Mecanismes Adaptatifs et Évolution UMR7179, CNRS, National Museum of Natural History, Paris, France

**Keywords:** ontogeny, mother-infant interactions, hand preference, hemispheric specialization, primates, *Papio anubis*

## Abstract

The most emblematic behavioral manifestation of human brain asymmetries is handedness. While the precise mechanisms behind the development of handedness are still widely debated, empirical evidences highlight that besides genetic factors, environmental factors may play a crucial role. As one of these factors, maternal cradling behavior may play a key role in the emergence of early handedness in the offspring. In the present study we followed 41 olive baboon (*Papio anubis*) infants living in different social groups with their mother for which direction (e.g., left- or right-arm) and degree of maternal cradling-side bias were available from our previous published study. We assessed hand preferences for an unimanual grasping task at 3 developmental stages: (1) 0-4, (2) 4-6 and (3) 9-10 months of age. We found that individual hand preferences for grasping exist as soon as the first months of age, with a population-level left-handedness predominance, being stable until 6 months; to wit the period during which juveniles are mainly carried by their mothers. More importantly, this early postnatal handedness is positively correlated with maternal cradling lateralization. Interestingly, hand preferences assessed later in the development, once juveniles are no longer carried (i.e., from 9 to 10 months of age), are less consistent with the earlier developmental stages and no longer dependent from the maternal cradling bias. Our findings suggest that the ontogenetic dynamics of the infant’s hand preference and its changes might ultimately rely on the degree of infant dependence from the mother across development.

**Research Highlights:**

- Early postnatal individual hand preferences are detected for unimanual food grasping within the first four months of age.
- Earliest measures of infant hand preference are positively correlated with measures of maternal cradling lateralization.
- Hand preferences assessed later in the development, from 9 to 10 months of age are less consistent with the earlier developmental stages and independent from maternal cradling bias.

## INTRODUCTION

The evolution of the human brain led to a cerebral lateralization allowing to effectively compensate for space constraints on the endocranial volume (Hofman et al., 2014). While some organs are duplicated (i.e., kidneys, lungs), the two hemispheres of the human brain display a functional specialization associated with structural asymmetries. This dissociation of specialized processes of left and right hemispheres allows to optimize the associated functions, for instance, the language for the left hemisphere, and emotions’ processing for the right one in a majority of individuals (Knecht et al., 2000; Gainotti, 2019). As the nerve fibers of the motor cortices are contralaterally innervated, the dominant hemisphere processes can manifest as contralateral motor behaviors (Hellige, 1993). Thus, the most emblematic behavioral manifestation of contralateral brain asymmetries is handedness. (Hammond, 2002)

The mechanisms that may influence the development of handedness are widely debated on both theoretical and empirical grounds. In humans, as hand preferences run in families, many studies have proposed genetic models (Annett, 1985; Laland et al., 1995; McManus & Bryden, 1992; Yeo & Gangestad, 1993), but no gene has been linked to the expression of handedness yet. In parallel, other studies have investigated nongenetic factors associated with the early developmental environment. As the mother acts on the immediate environment of the fetus and then of the infant (Damerose & Vauclair, 2002), these works namely focused on asymmetries in the prenatal environment and in the mother-infant interactions (Hopkins, 2004). Since the maternal intrauterine environment is asymmetric, it has been suggested that the position of the fetus may play a role in the development of lateralization in the motor system (Hopkins & Rönnqvist, 1998; Previc, 1991), such as handedness. Ultrasound scans have enabled to demonstrate that limb movements emerge during fetal life: young human fetuses already grab the umbilical cord, push the uterine wall, and even repeat hand-mouth contacts (Sparling et al., 1999; Takeshita et al., 2006). Those empirical evidence highlight that besides genetic factors, other nongenetic factors may play a role in the development of human handedness (Hopkins, 2004; Fagard, 2013).

Within this theoretical framework, non-human animal models have been used in comparative research for investigating factors influencing limb lateralization. While brain lateralization is not specific to humans but has been well established in many other vertebrates (see Rogers, 2002 for a review; Rogers & Andrew, 2002; Vallortigara & Bisazza, 2002; Vallortigara & Rogers, 2005), a large body of literature documents the lateralization of the primate forelimb motor system. In fact, in non-human primates, hand preference for various manual tasks has been reported at both individual- and population-levels (MacNeilage et al., 1987; Ward and Hopkins, 1993; Hopkins, 1996; Hook-Costigan & Rogers, 1997; Meguerditchian et al., 2012; Molesti et al., 2016). Interestingly, in chimpanzees and olive baboons specifically, the asymmetric use of the hands for complex manipulation activities, such as bimanual grasping, has been found, just like in humans, correlated to contralateral brain structural asymmetries within a section of the central sulcus related to the motor hand area (*Pan troglodytes*: Hopkins & Cantalupo, 2014; *Papio anubis*: Margiotoudi et al., 2019).

As mentioned above, a key factor of the developmental environment is the mother, acting on the immediate environment of the fetus and then of the infant (Damerose & Vauclair, 2002). In some of these primate species, asymmetries in mother-infant interactions have been documented, which makes non-human primates a relevant model to better understand the role of the early developmental environment in the emergence of asymmetric hand use. Indeed, in human and non-human primates such as chimpanzees, gorillas and baboons, maternal cradling of newborns is lateralized at individual-level and shows a left-side bias at population-level, which means the use of left arm is favored over the right arm to cradle the infant in a majority of individuals (Manning et al., 1994; Boulinguez-Ambroise et al., 2020). For instance, in chimpanzees, Hopkins et al. (1993) have found an inverse relationship between this maternal ventro-ventral cradling bias and the offspring hand preference for simple reaching at the age of 3 years. However, it is unclear whether the acquisition of handedness may be related to the infant’s hand that is free from the function of clinging, or to the hand used to hold onto the mother’s side, as they may receive different levels of motor and neurological stimulation (Hopkins, 2004). Early postnatal infant lateralization has been poorly investigated in non-human primates. The only few data available so far reported manual performance asymmetries in the strength of grasping responses in human and chimpanzee neonates (Petri & Peters, 1980; Fagot & Bard, 1995).

In the present study, we have investigated the development of hand preference in connection with side-asymmetries in maternal cradling in an Old-Word monkey: the olive baboon (*Papio anubis*). During the first months of life, the young baboon is manly cradled by the mother (Rose, 1977; Altmann & Samuels, 1992). The mother cradles her infant ventro-ventrally when sitting or even walking: she holds the infant supporting its weight with one arm, close to her body, in one of her peri-personal hemispaces. Maternal cradling is lateralized at both individual and population-levels in baboons (Boulinguez-Ambroise et al., 2020). If cradled on the left, the infant embraces and holds onto the left side of the mother with its right arm, the left hand being free, and vice versa. The hand that is not recruited for clinging on the fur, is free for reaching actions (towards conspecifics or objects), or even fine manipulative grasping actions (i.e., grasping food items on the ground, showing bouts of grooming). After 4 months, the frequency of infant-cradling decreases in favor of dorsal (i.e., riding) or ventral carrying (see Fig. 1), which requires from the infant the use of both hands to cling on the fur (Rose, 1977; Altmann & Samuels, 1992). After 9 months, infants are not carried by the mother anymore (Nash, 1978). According to maternal cradling side-asymmetries found in baboon mothers (dataset published by Boulinguez-Ambroise et al., 2020), we have tested the effect of this asymmetric mother-infant interaction on the development of the offspring handedness. We have thus longitudinally assessed the hand preferences for manual grasping tasks (i.e., unimanual food grasping) in 41 baboon infants following 3 different developmental stages, from 0 to 4 months, from 4 to 6 months, and from 9 to 10 months of age. The results will help us to determine whether or not: (1) baboons from a very young age show a manual lateralization for unimanual grasping, (2) maternal cradling-side bias influences those early patterns of infant manual lateralization, (3) individual hand preferences are stable across ontogeny.

**Figure 1.**
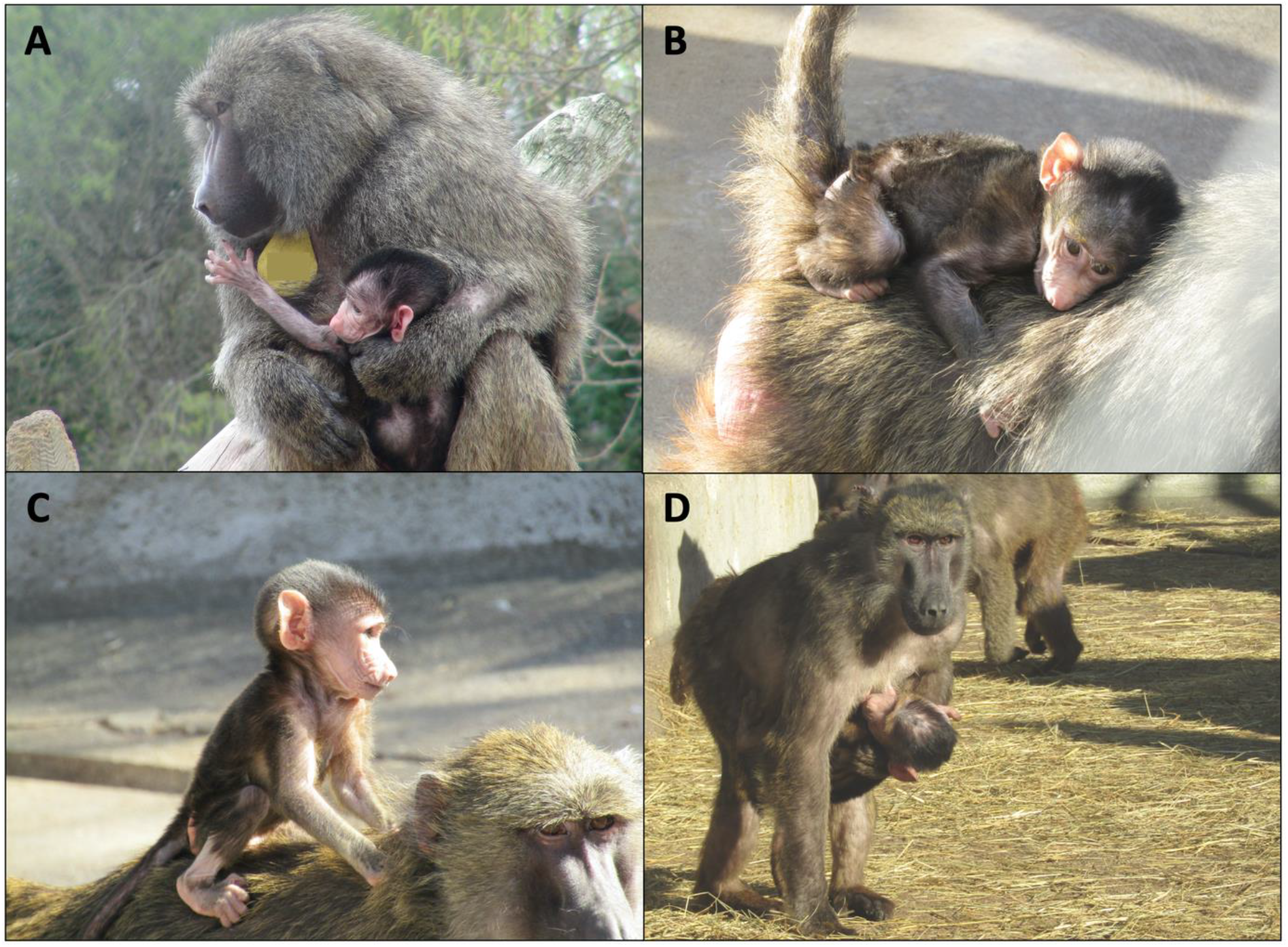
Pictures illustrating infant’s holding in female olive baboons. **A** The mother cradles the infant by supporting it with one arm. One of the infant’s arms embraces and holds onto the mother’s side, whereas the other arm is free. **B** The infant is carried dorsally, being lying down on her mother’s back. **C** The infant is carried dorsally, riding her mother’s back. **D** The mother carries her infant on her belly. The infant clings on her fur with both limbs, with or without a support by a mother’s arm. Photograph credit: Grégoire Boulinguez-Ambroise

## MATERIAL AND METHODS

### Subjects

All baboons (*Papio anubis*) were born and raised in captivity at the Primatology Station of the CNRS (UPS846 CNRS, Rousset-Sur-Arc, France, Agreement C130877). Subjects live in social groups, housed in large enriched aviaries or parks from 28 to 291 m^2^, containing multiple climbing structures. All enclosures comprise an outside area, as well as an inside sheltered space. In total, we assessed hand preference in 41 infant baboons (19 females and 22 males), aged from birth to 10 months old, from 30 mothers. We tested 33 young baboons (16 females, 17 males) for age 0 to 4 months, 29 subjects (13 females, 16 males) for age 4 to 6 months, and 25 subjects (11 females, 14 males) from age 9 to 10 months. For each of those 41 infants, the preferential use of their mother’s arm to cradle them (i.e., toward left *versus* right arm) was quantified in a previous study (available dataset published by Boulinguez-Ambroise et al., 2020).

### Ethics

The study was approved by the “C2EA-71 Ethical Committee of Neurosciences” (INT Marseille) under the number APAFIS#13553-201802151547729, and has been conducted at the Station de Primatologie (Rousset-Sur-Arc, France, Agreement C130877). All methods were performed in accordance with the relevant CNRS guidelines and the European Union regulations (Directive 2010/63/EU).

### Procedure of data collection

We assessed hand preference for unimanual food grasping at 3 different stages of the development of young olive baboons: 1) the first four months of life, period during which babies are mainly cradled by their mothers, one arm embracing and holding onto the side of the mother, the other arm being free 2) 4 to 6 months, period during which infants are less and less cradled but rather carried dorsally or ventrally, without a support by the mother’s arm, clinging on the fur with both limbs (Rose, 1977; Altmann & Samuels, 1992), and 3) after 9 months, as they are not carried anymore by the mother (Nash, 1978). Please, see figure 1 for illustrations of maternal infant’s holding. Subjects were tested in their social groups. According to a behavioral sampling procedure, we quantified the use of the left hand versus the right hand to grasp food items. Small grains and fruit cubes (i.e., 1cm^3^) were frequently scattered inside the enclosures in large quantities so as to maximize data collection and alleviate competition between individuals. We considered a food grasp as the following event: the subject spontaneously grasps a food item and brings it to its mouth. For a grasping occurrence to be counted, both of the subject’s hands had to be preliminary free (i.e., not holding an object in one hand), and the food had to be in front of the individual (i.e., not on its sides), please see figure 2. Thus, we assumed there was an independent choice of hands for grasping, with a limited influence of the posture. For each age range, a minimum of 30 grasping occurrences was required.

**Figure 2.**
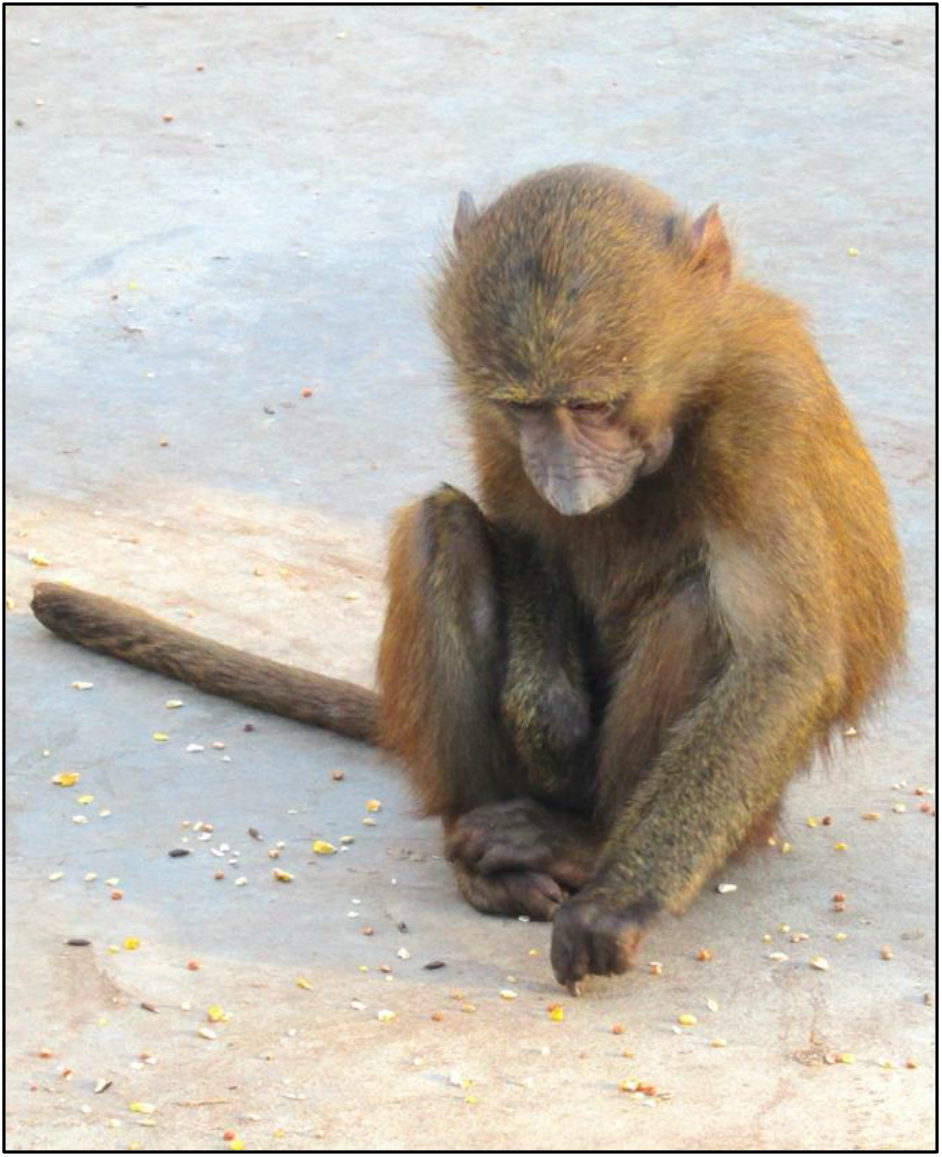
Pictures of juvenile olive baboons performing an unimanual grasping task. Photograph credit: Grégoire Boulinguez-Ambroise

### Statistical Analysis

For the three developmental stages (i.e., 0-4, 4-6, and 9 months of age), we determined the hand preference of each subject for unimanual grasping by calculating an individual Handedness Index (HI) using the formula (R−L)/(R+L). R and L represents the total right and left hand uses, respectively (Meguerditchian et al., 2011, 2013). A negative value indicates a left-hand preference, whereas a positive value a right-hand preference. The same formula was used by Boulinguez-Ambroise et al. (2020) to calculate the maternal cradling-side bias index score (CBI, based on the total left and right arm uses) of each juvenile we tested. The direction of cradling-side asymmetry was further determined for each infant by calculating a z-score, based on the total left and right arm uses (Meguerditchian et al., 2011), allowing to classify each mother as having a left side (z ≤ − 1.96) or right side (z ≥ 1.96) cradling bias, or behaving ambiguously (− 1.96 < z < 1.96). We first performed (1) Pearson correlation tests to assess whether hand preference was stable across the three development stages or not. We then performed the following statistical analyses for each developmental stage: we ran (2) a linear regression to test the effect of the maternal CBI as predictor of the infant’s HI. We further ran (3) a Kruskal–Wallis rank sum test with the cradling-side bias (i.e., based on z-score calculation) as a qualitative variable and the juvenile HI as a quantitative variable. (4) We also tested for a population-level left-handedness predominance by calculating the mean handedness index, and running a one-sample t test. We checked normality by performing a Shapiro–Wilk Normality test. We used the following RStudio packages: FactoMineR, car, MASS, readxl.

## RESULTS

### Development of asymmetric hand use for unimanual grasping

#### a) At individual-level

We found unimanual grasping to be strongly lateralized at the individual level in each of the three developmental stages (i.e., during the first 4 months after birth, between 4 and 6 months of age, from 9 to 10 months of age); see Table 1. According to a Pearson correlation, the individual Handedness Index (HI) were significantly correlated across the two first developmental stages, *r*_24_ = 0.6, *p* = 0.002, suggesting the stability of hand preference for unimanual grasping from 0 to 6 months. In contrast, between the third developmental stage and previous developmental stages, the correlation of HI remained positive but decreased and lost significance (between stages 3 and 1: *r*_18_= 0.39, *p* = 0.10; between stages 3 and 2: *r*_18_ = 0.36, *p* = 0.12, respectively), indicating that, after 6 months of age, patterns of individual hand preference change across time.

**Table 1.** Distribution of hand preference for an unimanual grasping task at three developmental stages in olive baboons (*Papio anubis*). Based on the calculation of z-scores, individuals are classified as being left-handed (z ≤ − 1.96), right-handed (z ≥ 1.96), or being ambiguously lateralized (− 1.96 < z < 1.96). The calculation of the z-score is based on the total left and right hand uses in an unimanual food grasping task. Mean.HI: Mean Handedness Index; SD: Standard Deviation. *p<0.10, **p<0.05

|  | 0-4 months<br>n=33 | 4-6 months<br>n=29 | 9-10 months<br>n=25 |
| --- | --- | --- | --- |
| Left-handed | 19 (58%) | 16 (55%) | 14 (56%) |
| Right-handed | 9 (27%) | 8 (28%) | 7 (28%) |
| Ambiguously handed | 5 (15 %) | 5 (17%) | 4 (16%) |
| Mean.HI ± SD | **-0.12 ± .39\*** | **-0.14 ± .36\*\*** | -0.07 ± .44 |

#### b) At population-level

During the first four months following birth, the calculation of the mean Handedness Index score among the total of 33 infants, Mean.HI = -0.12, SD = .39, showed a trend for a left-handedness predominance which approaches conventional level of significance according to an one-sample t-test (t_32_ = -1.78, *P* = 0.08). At the second developmental stage, from 4 to 6 months of age, the calculation of the mean Handedness Index score among the total of 29 infants, Mean.HI = -0.14, SD = .36, showed a significant left-handedness predominance according to a one-sample t-test (t_28_ = -2.13, *P* = 0.04). Around 9 months of age, the calculation of the mean Handedness Index score among the total of 25 infants, Mean.HI = -0.07, SD = .44, showed no population-level bias according to a one-sample t-test (t_24_ = -0.85, *P* = 0.41).

### Effect of maternal cradling asymmetries on the development of offspring hand preference (i.e., unimanual grasping)

#### a) During the first 4 months after birth

Linear model detected that the juvenile Handedness Index (HI) significantly increased with increasing Cradling Bias Index (F_1,31_ = 37.37; *P* < 0.0001). By running a Kruskal-Wallis rank sum test with the cradling-side bias (i.e., based on z-score calculation) as a qualitative variable and the juvenile HI as a quantitative variable, we found that the females that cradle their infants on the left side have offspring with significant lower HI (Mean.HI = -0.34 < 0) than the right cradling mothers (Mean.HI = 0.33 > 0; Kruskal-Wallis *X*^*2*^ = 13.8, *P* < 0.001; see Fig.3). Regarding the 5 ambiguously lateralized cases of cradling, three infants were ambidextrous, one was left-handed and one was right-handed (based on z-score calculation).

**Figure 3.**
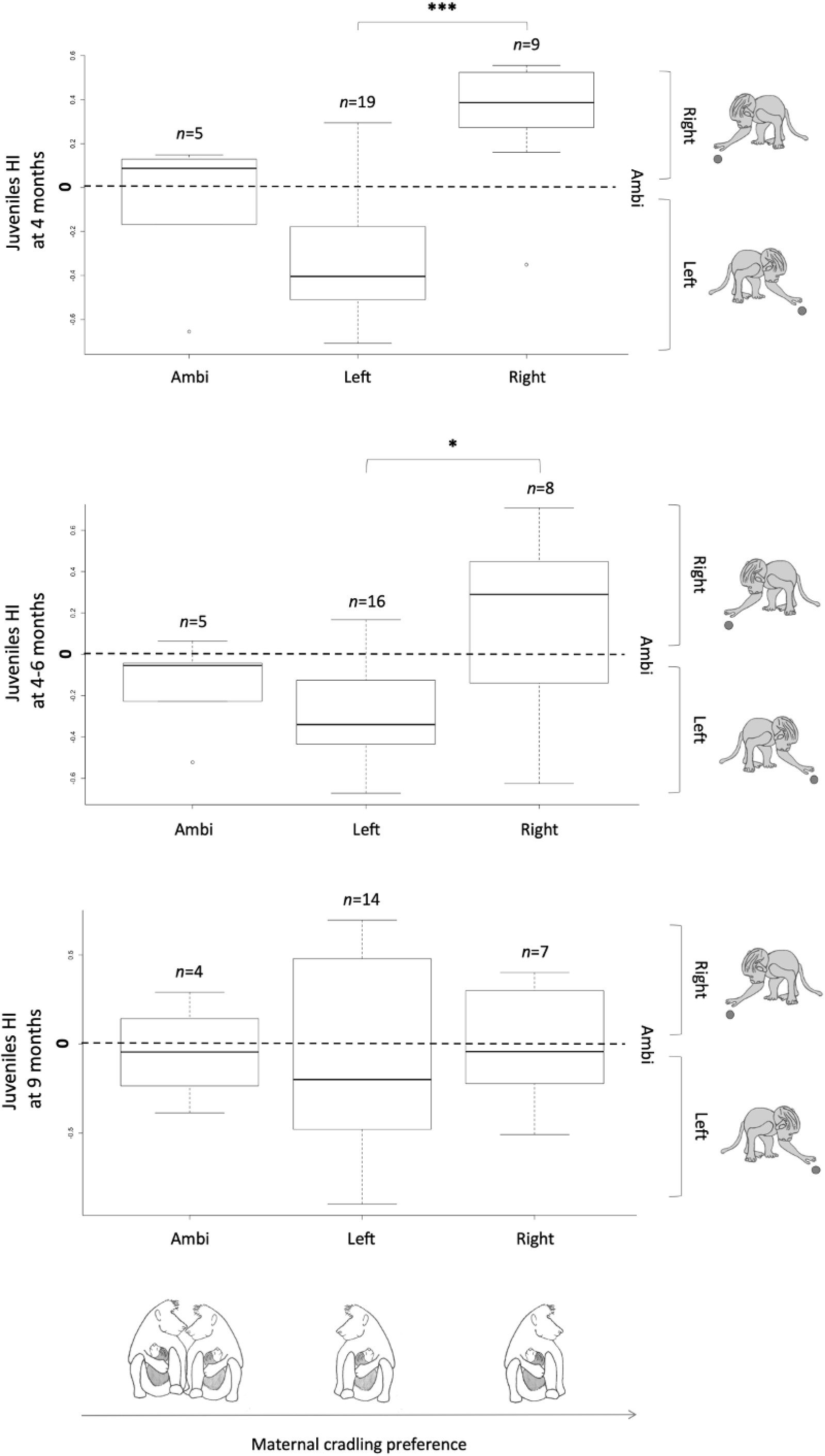
Effect of maternal cradling bias on the development of hand use in juvenile olive baboons (*Papio anubis*). Boxplot of the Handedness Index (HI) of juveniles according to the maternal cradling-side bias, (**A**) during their first four months of life (n=33), (**B**) between 4 and 6 months of age (n=27), and (**C**) around 9 months of age (n=25). The calculation of the Handedness Index (HI) is based on the total left and right arm uses; a negative value indicates a left-hand preference, whereas a positive value a right-hand preference. The maternal cradling-side bias was assessed by Boulinguez-Ambroise et al. (2020). (*P*-value * < 0.05; *** < 0.001). The boxplots are made of a vector containing the 1st quartile (Q1, box lower ‘hinge’), the median (bold horizontal line), the 3rd quartile (Q3, box upper ‘hinge’), and the adjacent values (whiskers). The length of the whiskers is calculated from the interquartile range (IQR = Q3 - Q1): Q1 -1.5*IQR (lower whisker), Q3 + 1.5*IQR (upper whisker).

#### b) Between 4 and 6 months of age

Linear model detected that the offspring Handedness Index is positively related to the maternal Cradling Bias Index between 4 and 6 months of age (F_1,27_ = 14.18; P < 0.001; see Fig.3). By running a Kruskal-Wallis rank sum test with the cradling-side bias (i.e., based on z-score calculation) as a qualitative variable and the juvenile HI as a quantitative variable, we found that the females that cradle their infants on the left side have offspring with significant lower HI (mean HI = -0.29 < 0) than the right cradling mothers (Mean.HI = 0.16 > 0; Kruskal-Wallis *X*^*2*^ = 4.59, *P* = 0.032; see Fig.3). Regarding the 5 ambiguously lateralized cases of cradling, three infants were ambidextrous, and two were left-handed (based on z-score calculation).

#### c) Around 9 months of age

Handedness Index of 9-months old juveniles was not related to the maternal Cradling Bias Index (F_1,23_ = 0.88; *P* = 0.36; see Fig.3).

## DISCUSSION

The present study provides several straightforward findings. First, our results showed early postnatal individual hand preference for unimanual food grasping within the first four months of age (i.e., 1^st^ stage). Second, we found that, while early individual hand preferences were stable until 6 months of age (i.e., 2^nd^ stage), the hand preferences assessed later in the development, from 9 to 10 months of age (i.e., 3^rd^stage), were less consistent with the earlier developmental stages. Third, at the population-level, we reported a significant left-handedness predominance for unimanual grasping until 6 months of age (i.e., 1^st^ and 2^nd^ stages), while the bias toward left-handedness is not significant anymore from 9 months (i.e., 3rd stage). Fourth, we found that the measures of infant hand preference (i.e., Handedness Index: HI) were significantly correlated with the measures of maternal cradling lateralization (i.e., Cradling Bias Index: CBI) at the first stage of development (i.e., 0-4 months of age). Most infants that were cradled on the left arm of their mother showed a left-hand preference for grasping (i.e., negative value of HI), whereas infants that were cradled on the right arm, a right-hand preference (i.e., positive value of HI). Then, the relation between the asymmetry of maternal cradling and the infant hand preference is less pronounced at the 2nd stage (i.e., 4-6 months of age), and was no longer present later in the last developmental stage (i.e., 9-10 months of age).

One potential interpretation is that the ontogenetic dynamics of the infant’s hand preference and its changes might ultimately rely on the degree of infant dependence from the mother across development.

At the first stage, the infants are mainly cradled by their mothers (Rose, 1977; Altmann & Samuels, 1992) and the effect of maternal cradling side bias on the offspring’s hand preference is maximal. We suggest the maternal asymmetric cradling behavior, expressed during this earliest age class on the infant, has a direct impact on how the infant develops hand preference when experiencing object grasping outside its mother. Specifically, maternal cradling would favour the development of the infant’s hand preference for the hand free from the function of clinging. For instance, when cradled on the left side (and vice versa when cradled on the right side), it is the infant’s right hand which falls on the mother’s side and is thus recruited for clinging or holding on her. In contrast, the infant’s left hand could be free to interact with the external environment including reaching actions towards objects (e.g., fine manipulative food grasping actions) or towards conspecifics (e.g., grooming) involving thus greater motor and neurological stimulation than the other hand, inducing ultimately early development of hand use for food grasping when the infant is behaving outside the mother’s embrace. Moreover, we found that functional manual asymmetry for grasping was actually not present in ambiguously cradled infants.

At the 2nd stage of development, we noticed an alteration of the cradling side effect on infant hand preference for grasping. It might be explained by the decrease of frequencies of asymmetric maternal cradling in favor of middle dorsal or ventral infant carrying, (Rose, 1977; Altmann & Samuels, 1992), and of independent locomotion. Nevertheless, such an alteration of the effect seems to not be strong enough to impact patterns of infant’s hand preference across the two first age classes (i.e., 1^st^and 2^nd^ stages, until 6 months of age) which remain consistent at both the individual - and population-levels. Infants are predominantly left-handed forgrasping at these two early developmental stages, which could be attributed to the maternal cradling effect. In fact, as we found in a previous study, olive baboon mothers exhibit a left-cradling bias at population-level (Boulinguez-Ambroise et al., 2020), which ultimately impacted the leftward direction of the population-level offspring handedness. Maternal left-cradling bias has been also demonstrated in great apes (chimpanzees and gorillas; Manning et al., 1994) and humans (Malatesta et al., 2019). Such phenomenon seems not to be related with the mother handedness (Forrester et al., 2019), but would rather reflect the right-hemispheric dominance for emotional processing (Manning & Chamberlain, 1991). In fact, the brain right hemisphere is specialized in the perception of emotional facial expressions (Bryden & Levy, 1983; Gainotti, 2019). Since left-side cradling exposes the baby face to the left visual field of the mother, which is projected mainly to her right brain hemisphere, this would favour the mother’s monitoring of the emotional state of the infant. According to our present findings, it might be thus not excluded that it is the brain specialization for emotion processing of the mother which has an indirect influence - though asymmetric cradling behavior - on the development of hand use in infant baboons. In chimpanzees, an inverse relationship was found between the maternal cradling bias and the offspring hand preference (Hopkins et al., 1993); but the authors tested juveniles for simple reaching at a much later developmental stage (i.e., 3 years of age).

Finally, the loss of the maternal cradling side’s effect on the offspring’s hand preference at the 3rd stage of development might be due to the complete autonomy of offspring which is usually weaned from the mother after 9 months (Nash, 1978). This is consistent with the changes we noticed in the pattern of infants’ hand preference at the third stage of development at both individual- and population-levels (i.e., inconsistency with earlier age classes and loss of population-level left-handedness). This lack of population-level handedness later in development is consistent with what was found previously in adult baboons, namely significant individual hand preference for unimanual grasping but no significant population-level handedness (Vauclair et al., 2005; Molesti et al., 2016). Whether the pattern of individual handedness for unimanual grasping will keep changing or remain stable after 9-10 months until adulthood, is unclear. In a previous study in olive baboons, Molesti et al. (2016) found that individual patterns of hand preference for unimanual grasping remain quite consistent across time in adults.

Further questions would address the ontogenesis of handedness at latter stages across primate’s development (i.e., above 10 months of age) following the emergence of manual behaviors with higher motor demand than grasping. For instance, follow-up study in this cohort of young baboons would help determine the potential role of the emergence of “bimanual coordinated manipulations” on handedness’ ontogeny. Such behaviors (referred as “bimanual tasks” in the rest of the paper), which consist of using the two hands in an asymmetric but complementary matter (e.g., holding a tube with one hand and removing the food inside a tube with the other hand), have received special attention and interest. In humans, while evidence of manual lateralization has been reported even before birth (Hepper et al., 1991) and for unimanual behaviors at around 5 and 6 months of age (Ramsay, 1980; Michel, 1981, 1982; Michel et al. 1985), the appearance of a reliable hand-use preference for role-differentiated bimanual manipulations were documented around the age of 13 months (Kimmerle et al., 2010).

In fact, even if in both human children and olive baboons, the direction of individual hand preference was found quite consistent between unimanual and bimanual tasks (Molesti et al., 2016; Fagard & Marks, 2000), bimanual tasks were proposed to constitute better measures of hand preference for the following reasons. In comparison to unimanual grasping, bimanual tasks elicited (1) hand use measures with lower sensitiveness to situational confounding factors such as the subjects’ posture or the initial position of the item to reach, (2) higher motor demands which increase the probability of using of the preferred hand. Therefore, in both children (Fagard & Marks, 2000; Potier et al., 2013) and nonhuman primates (Meguerditchian et al., 2013 for a review) such as baboons *Papio anubis* (Vauclair, 2005; Molesti et al., 2016), bimanual tasks were found to elicit not only stronger right-handedness predominance at the population-level but also more robust individual hand preferences in terms of degree/strength of manual asymmetry and consistency across time. Interestingly, direction and degree of hand preference for a bimanual task in nonhuman primates such as baboons, capuchin monkeys, or chimpanzees have been found to be associated with contralateral neuro-structural asymmetries in the primary motor cortex including the surface of the motor hand area surface, its neuronal densities or its adjacent Central sulcus depth (Hopkins and Cantalupo, 2004; Dadda et al., 2006; Sherwood et al., 2007; Phillips & Sherwood, 2005; Hopkins, 2013; Margiotoudi et al., 2019).

## CONCLUSION

The results of this study show a clear effect of a nongenetic factor on the development of hand use in young olive baboons. The infant handedness is strongly related to the maternal cradling side bias during the first 6 months of life; to wit the period during which the juveniles are mainly carried by their mothers. Once juveniles are no longer carried (i.e., at 9 months), the effect of this mother-infant interaction on infant handedness is no longer present, and hand preference can even be reversed. Follow-up investigations on this cohort of growing baboons would help us to determine at which age of the development the hand preferences for unimanual grasping start to (1) stabilize across time, (2) predict the hand preference for bimanual coordinated actions, or even - with the contribution of MRI imaging approach - (3), predict contralateral hemispheric specialization of the brain within the central sulcus and its potential change across ontogeny.

## ACKNOWLEDGMENTS

We are particularly thankful to Jérémy Roche, Théophane Piette and Léa Langérôme for their assistance in data collection as well as Siham Bouziane, Solène Bischoff & Solène Dulac. We warmly thank Pascaline Boitelle (veterinarian) and the animal keepers of the CNRS UPS 846. We thank the Center for Research and Interdisciplinarity (CRI, Paris) for financial aid. The project has received funding from the European Research Council under the European Union’s Horizon 2020 research and innovation program Grant Agreement No. 716931—GESTIMAGE—ERC-2016-STG (P.I. Adrien Meguerditchian), as well as from the French “Agence Nationale de la Recherche” ANR-16-CONV-0002 (ILCB) and the Excellence Initiative of Aix-Marseille University (A*MIDEX).

## REFERENCES

Altmann, J. & Samuels, A. 1992. Costs of maternal care: Infant-carrying in baboons. Behav. Ecol. Sociobiol., 29, 391–398.

Annett, M. 1985. Left, Right, Hand, and Brain: The Right-Shift Theory, Erlbaum, London.

Bellugi, U. 1991. The link between hand and brain: Implications from a visual language. In D. S. Martin (Ed.), Advances in cognition, education, and deafness. Washington, DC: Gallaudet Univ. Press. 11–35.

Boulinguez-Ambroise, G., Pouydebat, E., Disarbois, É. & Meguerditchian, A. 2020. Human-like maternal left-cradling bias in monkeys is altered by social pressure. Sci. Rep., 10, 11036.

Bryden, M. P. & Levy, R. G. 1983. In Neuropsychology of Human Emotion, eds K.M. Heilman & P. Satz (Guilford Press, New York).

Dadda, M., Cantalupo, C., Hopkins, W.D. 2006. Further evidence of an association between handedness and neuroanatomical asymmetries in the primary motor cortex of chimpanzees (Pan troglodytes). Neuropsychologia, 44(12), 2582–2586.

Damerose, E. & Vauclair, J. 2002. Posture and laterality in human and non-human primates: Asymmetries in maternal handling and infant’s early motor asymmetries. In Rogers, L., and Andrew, R. J. (eds.). Comparative Vertebrate Lateralization, Oxford University Press, Oxford, pp. 306–362.

Erwin, J., Anderson, B. & Bunger, D. 1975. Nursing behavior of infant pigtail monkeys (Macaca nemestrina): Preferences for nipples. Percept. Mot. Skills 592–594.

Fagot, J. & Bard, K.A. 1995. Asymmetric grasping response in neonate chimpanzees (Pan troglodytes). Infant Behav. Dev., 18, 253–255.

Forrester, G.S., Davis, R., Mareschal, D., Malatesta, G. & Todd, B.K. 2019. The left cradling bias: an evolutionary facilitator of social cognition? Cortex, 118, 116–131.

Fagard, J. & Marks, A. 2000. Unimanual and bimanual tasks and the assessment of handedness in toddlers. Dev. Sci., 3, 137–147.

Fagard, J. 2013. The nature and nurture of human infant hand preference. Ann N Y Acad Sci., 1288, 114–23.

Gainotti, G. 2019. Emotions and the right hemisphere: can new data clarify old models? Neuroscientist, 25, 258–270.

Hammond, G. 2002. Correlates of human handedness in primary motor cortex: A review and hypothesis. Neurosci. Biobehav. Rev., 26, 285–292.

Hellige, J.B. 1993. Unity of thought and action: Varieties of interaction between the left and right cerebral hemispheres. Curr. Dir. Psychol. Sci., 2(1), 21–26.

Hepper, P.G., Shahidullah, S. & White, R. 1991. Handedness in the human fetus. Neuropsychologia, 29, 1107–1111.

Hofman, M. A. 2014. Evolution of the human brain: when bigger is better. Front. Neuroanat., 8, 15.

Hook-Costigan, M.A. & Rogers, L.J. 1997. Hand preferences in New World primates. Int. J. Comp. Psychol., 9, 173–207.

Hopkins, W.D., Bard, K.A., Jones, A. & Bales, S. 1993. Chimpanzee hand preference for throwing and infant cradling: Implications for the origin of human handedness. Curr. Anthropol., 34, 786–790.

Hopkins, W.D. & Bard, K.A. 1995. Evidence of asymmetries in spontaneous head turning in infant chimpanzees (Pan troglodytes). Behav. Neurosci., 109, 808–812.

Hopkins, W.D. 1996. Chimpanzee handedness revisited: 54 years since Finch (1941). Psychonom. Bull. Rev., 3, 449–457.

Hopkins, B. & Rönnqvist, L. 1998. Human handedness: Developmental and evolutionary perspectives. In Simon, F., and Butterworth, G. (eds.), The Development of Sensory, Motor and Cognitive Capacities of Early Infancy: From Perception to Cognition, Psychology Press, East Sussex, UK, pp. 191–236.

Hopkins, W.D. & Bard, K.A. 2000. A longitudinal study of hand preference in chimpanzees (Pan troglodytes). Dev. Psychobiol., 34, 292–300.

Hopkins, W.D. 2004. Laterality in Maternal Cradling and Infant Positional Biases: Implications for the Development and Evolution of Hand Preferences in Nonhuman Primates. Int. J. Primatol., 25, 1243–1265.

Hopkins, W.D. & Cantalupo, C. 2004. Handedness in chimpanzees (Pan troglodytes) is associated with asymmetries of the primary motor cortex but not with homologous language areas. Behav. Neurosci., 118(6), 1176.

Hopkins, W.D. 2013. Neuroanatomical asymmetries and handedness in chimpanzees (Pan troglodytes): a case for continuity in the evolution of hemispheric specialization. Ann. N. Y. Acad. Sci., 1288(1), 17–35.

Kimmerle M., Ferre, C.L., Kotwica, K.A. & Michel, G.F. 2010. Development of role-differentiated bimanual manipulation during the infant’s first year. Dev. Psychobiol., 52, 168–80.

Knecht, S., Dräger, B., Deppe, M., Bobe, L., Lohmann, H., Flöel, A., et al. 2000. Handedness and hemispheric language dominance in healthy humans. Brain, 123(12), 2512–2518.

Laland, K.N., Kumm, J., Van Horn, J.D. & Feldman, M.W. 1995. A gene-culture model of human handedness. Behav. Genet., 25, 433–445.

Levitt, P. 2003. Structural and functional maturation of th e developing primate brain. J. Pediatr., 143, 35–45.

MacNeilage, P.F., Studdert-Kennedy, M.G. & Lindblom, B. 1987. Primate handedness reconsidered. Behav. Brain Sci., 10, 247–303.

Malatesta, G., Marzoli, D., Rapino, M. & Tommasi, L. 2019. The left-cradling bias and its relationship with empathy and depression. Sci. Rep., 9, 6141.

Manning, J.T. & Chamberlain, A.T. 1991. Left-side cradling and brain lateralization. Ethol. Sociobiol., 12, 237–244.

Manning, J.T., Heaton, R. & Chamberlain, A.T. 1994. Left-side cradling: Similarities and differences between apes and humans. J. Hum. Evol., 26, 77–83.

Margiotoudi, K., Marie, D., Claidière, N., Coulon, O., Roth, M., Nazarian, B., Lacoste, R., Hopkins, W.D., Molesti, S., Fresnais, P., Anton, J.-L. & Meguerditchian, A. 2019. Handedness in monkeys reflects hemispheric specialization within the central sulcus. An in vivo MRI study in right- and left-handed olive baboons. Cortex, 118, 203–211.

McManus, I.C. & Bryden, M.P. 1992. The genetics of handedness, cerebral dominance and lateralization. In Rapin, I., and Segalowitz, S.J. (eds.), Handbook of Neuropsychology. Vol 6. Developmental Neuropsychology, Part 1, Elsevier, Amsterdam, pp. 115–144.

Meguerditchian, A., Molesti, S. & Vauclair, J. 2011. Right-handedness predominance in 162 baboons (Papio anubis) for gestural communication: Consistency across time and groups. Behav. Neurosci., 125, 653–660.

Meguerditchian, A., Donnot, J., Molesti, S., Francioly, R. & Vauclair, J. 2012. Sex difference in squirrel monkeys’ handedness for unimanual and bimanual coordinated tasks. Anim. Behav., 83, 635–643.

Meguerditchian, A., Vauclair, J. & Hopkins, W. D. 2013. On the origins of human handedness and language: A comparative review of hand preferences for bimanual coordinated actions and gestural communication in nonhuman primates: On the Origins of Human Handedness and Language. Dev. Psychobiol., 55, 637–650.

Meguerditchian, A., Phillips, K.A., Chapelain, A., Mahovetz, L.M., Milne, S., Stoinski, T., Bania, A., Lonsdorf, E., Schaeffer, J., Russell, J. & Hopkins, W.D. 2015. Handedness for unimanual grasping in 564 great apes: The effect on grip morphology and a comparison with hand use for a bimanual coordinated task. Front. Psychol., 6, 1794.

Michel, G.F. 1981. Right-handedness: A consequence of infant supine head-orientation preference? Science, 212, 685–687.

Michel, G.F. 1982. Ontogenetic precursors of infant handedness. Infant Behav. Dev., 5, 156.

Michel, G.F., Ovrut, M.R. & Harkins, D.A. 1985. Hand-use preference for reaching and object manipulation in 6-through 13-month-old infants. Genet. Soc. Gen. Psychol. Monogr., 111, 409–427.

Molesti, S., Vauclair, J. & Meguerditchian, A. 2016. Hand preferences for unimanual and bimanual coordinated actions in olive baboons (Papio anubis): consistency over time and across populations. J. Comp. Psychol., 130, 341–350.

Nash, L.T. 1978. The development of the mother-infant relationship in wild baboons (Papio anubis). Anim. Behav., 26(3), 746–59.

Nishida, T. 1993. Left nipple suckling preference in wild chimpanzees. Ethol. Sociobiol., 14. 45–52.

Petrie, B.F. & Peters, M. 1980. Handedness: Left/right differences in intensity of grasp response and duration of rattle holding in infants. Infant Behav. Dev., 3, 215–221.

Phillips, K.A., Sherwood, C.C. 2005. Primary motor cortex asymmetry is correlated with handedness in capuchin monkeys (Cebus apella). Behav. Neurosci., 119(6), 1701.

Potier, C., Meguerditchian, A., Fagard, J. 2013. Handedness for bimanual coordinated actions in infants as a function of grip morphology. Laterality, 18(5), 576–93.

Previc, F.H. 1991. A general theory concerning the prenatal origins of cerebral lateralization in humans. Psychol. Rev., 98, 299–334.

Ramsay, D.S. 1980. Onset of unimanual handedness in infants. Infant Behav. Dev., 3, 377–385.

Rogers, L.J. 2002. Lateralization in vertebrates: Its early evolution, general pattern, and development. In Slater, P. J. B., Rosenblatt, J. S., Snowdon, C. T., and Roper, T. J. (eds), Advances in the Study of Behavior, Vol. 31, Academic Press, San Diego, pp. 107–161.

Rogers, L.J. & Andrew, R.J. 2002. Comparative Vertebrate Lateralization, Cambridge University Press, Cambridge, UK.

Rose, M. 1977. Positional behavior of olive baboons (Papio anubis) and its relationship to maintenance and social activities. Primates, 18, 59–116.

Sherwood, C.C., Wahl, E., Erwin, J.M., Hof, P.R., Hopkins, W.D. 2007. Histological asymmetries of primary motor cortex predict handedness in chimpanzees (Pan troglodytes). J. Comp. Neurol., 503, 525–537.

Sparling, J.W, Van Tol, J. & Chescheir, N.C. 1999. Fetal and neonatal hand movement. Phys. Therapy, 79, 24–39.

Takeshita H., Myowa-Yamakoshi M. & Hirata S. 2006. A new comparative perspective on prenatal motor behaviors: Preliminary research with four-dimensional ultrasonography. In: Matsuzawa T., Tomonaga M., Tanaka M. (eds) Cognitive development in chimpanzees. Springer, Tokyo.

Tomaszycki, M., Cline, C., Griffin, B., Maestripieri, D. & Hopkins, W.D. 1998. Maternal cradling and infant nipple preferences in rhesus monkeys (Macaca mulatta). Dev. Psychobiol., 32, 305–312.

Vallortigara, G. & Bisazza, A. 2002. How ancient is brain lateralization? In Rogers, L. J., and Andrew, J. R. (eds.), Comparative Vertebrate Lateralization, Cambridge University Press, Cambridge, UK, pp. 9–69.

Vallortigara, G. & Rogers, L. J. 2005. Survival with an asymmetrical brain: advantages and disadvantages of cerebral lateralization. Behav. Brain Sci., 28, 575–633.

Vauclair, J., Meguerditchian, A. & Hopkins, W.D. 2005. Hand preferences for unimanual and coordinated bimanual tasks in baboons (Papio anubis). Cogn. Brain Res., 25, 210–216.

Ward, J.P. & Hopkins, W.D. 1993. Primate Laterality: Current Behavioral Evidence of Primate Asymmetries, Springer-Verlag, New York.

Westergaard, G.G., Byrne, G. & Suomi, S.J. 1998. Early lateral bias in tufted capuchins (Cebus apella). Dev. Psychobiol., 32, 45–50.

Yeo, R. & Gangsted, S.W. 1993. Developmental origins of variation in human hand preference. Genetica, 89, 281–296.

